# Decoding social knowledge in the human brain

**DOI:** 10.1101/2020.08.18.255513

**Authors:** Daniel Alcalá-López, David Soto

**Affiliations:** Basque Center on Cognition, Brain and Language, San Sebastian, Spain; Ikerbasque, Basque Foundation for Science, Bilbao, Spain

## Abstract

The present functional MRI study addressed how the brain maps different aspects of social information. We focused on two key dimensions of social knowledge: affect and likableness. Thirty participants were presented with audio definitions, half referring to affective (e.g. *empathetic*) and half to non-affective concepts (e.g. *intelligent*). Orthogonally, half of the concepts were highly likable (e.g. *sincere*) and half were socially undesirable (e.g. *liar*). We used a support vector machine to delineate how both concept dimensions are represented in a set of 9 *a priori* brain regions defined from previous meta-analyses on semantic and social cognition. We show that average decoding in semantic regions (e.g. lateral temporal lobe, inferior frontal gyrus, and precuneus) outperformed social ROIs (e.g. insula and anterior cingulate), with the lateral temporal lobe containing the highest amount of information about the affect and likableness of social concepts. We also found that the insula had a bias towards affect while the likableness dimension was better represented in anterior cingulate cortex. Our results do not support a modular view of social knowledge representation. They rather indicate that the brain representation of social concepts implicates a distributed network of regions that involves ‘domain-specific’ social cognitive systems, but with a greater dependence on language-semantic processing.

## 1 Introduction

The past two decades have witnessed a flourishing interest in the study of brain representations of meaning (Bauer and Just 2019; Martin 2007), with a growing body of research outlining the mechanisms by which concrete concepts (e.g. *cat* or *scissors*) are represented in the brain. Early functional MRI studies (Haxby et al. 2001; Mitchell, Hutchinson, Just, et al. 2003) showed that brain activity patterns in a set of brain regions, usually referred to as the “semantic network”, carry rich information about the concept that the observer is experiencing (e.g. *animals* vs. *tools*). This network of brain regions involved in semantic processing comprises areas in the temporal, inferior frontal and parietal lobes, the medial prefrontal cortex, alongside the superior and inferior temporal lobes (Binder, Desai, et al. 2009). These findings have sparked a lively discussion on the nature of meaning representations. Originally, theoretical models suggested that the brain represents concepts as amodal symbols (Fodor 1975, 1998). A more recent approach argues that conceptual representations are grounded in the sensorimotor experiences associated with them (Barsalou 1999; Prinz 2004), which can be re-enacted via mental simulation processes (Soto et al. 2020). This grounded cognition framework was initially conceived the study of concrete concepts. Only recently there has been a similar attempt at studying the representations of abstract concepts. Unlike concrete concepts, these are not perceptually bound to a physical object as referent, which points in the direction of grounding abstract concepts beyond pure sensorimotor systems (Shea 2018). Instead, abstract concepts are probably grounded in more complex representations that involve events or situations that can only rely on perceptual and action systems to a certain extent (Wilson-Mendenhall et al. 2011). Put simply, domain-specific systems appear insufficient to allow us to distinguish closely related concepts such as *agreeable* and *joyful*. This view is congruent with a recent two-systems proposal of meaning representation by Borghi and cols. (2019). These authors argue that, although a sensorimotor feature-based system would be common for the representation of both concrete and abstract concepts, the latter would need the assistance of an additional system that incorporates more complex linguistic and social information (Borghi et al. 2019).

Here, we used functional MRI and multivariate pattern analysis (MVPA) to address how the human brain represents abstract social knowledge. Studies using mass-univariate approaches in fMRI data have pointed to brain areas showing partially overlapping activation during the presentation of abstract and concrete concepts, including key areas of the semantic network such as inferior frontal, premotor, and superior temporal areas (Binder, Westbury, et al. 2005). However, more recent attempts to study the neural representations of abstract concepts using MVPA have shown that higher-order regions in medial frontal areas, which are less related to sensory processing (Margulies et al. 2016), are particularly involved in the representation of abstract–relative to concrete–concepts. For instance, Ghio and others (Ghio et al. 2016) showed in an fMRI study that brain activity patterns in the inferior frontal gyrus and the insula allowed them to classify abstract vs. concrete concepts (e.g. emotional or mathematical vs. action concepts). This study further highlighted that fine-grained representations of conceptual categories appear to co-exist along the concrete-to-abstract continuum (e.g. number, emotion, moral, aesthetic or social concepts). Critically, social concepts differ from other (non-social) abstract concepts in that their social character is heavily influenced by contextual information. For instance, a concept such as *trustworthy* can relate to ourselves, to others, as well as to non-human animals or even inanimate objects.

Despite this promising research, the study of how the human brain implements the representation of social knowledge is still at an early stage of development. Perhaps one of the more robust findings indicates that the anterior temporal lobe (ATL) plays a key role in the processing of socially-relevant information. Many studies have found activity increases in this brain region when participants are presented with different sources of information regarding other individuals (Binney et al. 2016; Lin et al. 2018; Pobric et al. 2016; Zahn et al. 2007). Crucially, most of these studies used standard mass-univariate models of voxel activation. These approaches are not best-suited for assessing the brain basis of abstract concept representation because they lack the resolution to pinpoint meaningful representations of different types of social concepts that may be encoded in the multi-voxel patterns of fMRI activity. A recent fMRI study combined both uni- and multivariate approaches to study the neural representations of social concepts. The authors asked participants to learn biographical details of fictitious characters (Y. Wang et al. 2017). In a memory retrieval task, participants were cued with a subset of the study items before answering questions about a specific character. They found that the ATL serves as an integration hub for a distributed neural circuit that represents information related to people.

The medial prefrontal cortex (MFPC) has also been implicated in the representation of psychological traits. In an fMRI study, Ma and cols. (2014) asked participants to infer the psychological traits of other individuals from a series of descriptions. First, sentence with an implicit psychological trait was presented. Then, a second target sentence appeared and participants had to infer the individual’s trait, which could be either congruent or not with the first sentence. They found that activity patterns within the ventral MPFC showed a neural adaptation effect indicative of the representation of trait knowledge: when the target psychological trait was congruent with the prior implicit sentence, neural activation decreased faster than during incongruent psychological trait descriptions (Ma et al. 2014). It is clear that social information is composed of a range of underlying dimensions, yet it is unknown whether and how the brain maps seemingly different aspects of social information. A fundamental, low-dimensional feature underlying many psychosocial phenomena is affect (Barrett and Bliss-Moreau 2009). There is general agreement that affect involves a bodily state of certain valence and arousal (Barrett 2016). In the present study, we selected 18 concepts whose definition made an explicit mention to the affective state of people, whereas the remaining 18 concept definition did not include any direct reference to affect. On a different note, a large tradition of psychological research has emphasized the extent to which human judgements of one’s own and others’ experiences are deeply influenced by their social desirability or likableness (Anderson 1968; Fisher et al. 1985; Norman 1967). This research suggests that whether certain behaviors are likable or appropriate might be an underlying feature of the representation of interpersonal behaviors. Nevertheless, this topic has not been addressed in the neurosemantics literature.

Accordingly, here we investigated the brain representations of these abstract concepts associated with two fundamental processes in social cognition. On the one hand, we analyzed how the emotional content of social concepts is represented in the brain by contrasting concepts related to the affective states of other people, such as *cruel* or *caring*, and concepts that refer to the (non-affective) mental states of other people, such as *honest* or *intelligent*. On the other hand, the present study addressed how the likableness of social concepts is represented in the brain by comparing highly likable concepts, such as *empathetic* or *understanding*, to highly unlikable concepts, such as *phony* or *insensitive*.

## 2 Methods

### 2.1 Participants

We scanned thirty participants (mean age 24.07 *±* 3.67 years; 18 females). All of them had normal or corrected-to-normal vision, gave written informed consent prior to the experiment, and were financially compensated with 20 euros for their participation. The experiment lasted for about an hour and a half and was approved by the BCBL Ethics Review Board in compliance with the Helsinki Declaration.

### 2.2 MRI acquisition

The present fMRI study was performed on a SIEMENS’s Magnetom Prisma-fit scanner with a 3T magnet and a 64-channel head coil. We collected one high-resolution T1-weighted image and eight functional runs for each participant. Each functional run consisted of a multiband gradient echo-planar imaging sequence with an acceleration factor of 6, resolution of 2.4 × 2.4 × 2.4mm^3^, TR of 850 ms, TE of 35 ms, and a bandwidth of 2582Hz/Px was used to obtain 537 3D volumes of the whole brain (66 slices; FOV = 210 mm).

The auditory stimuli for the experimental task (i.e. the concept definitions) were presented through insert earphones (S14, Sensimetrics, Malden, MA). Presentation volume was adjusted to a comfortable level for each participant. The visual elements of the experimental setup (e.g. fixation cross and stop signal) were projected on an MRI-compatible, out-of-bore screen using a projector in the adjacent room. A microphone mounted in the head coil allowed participants to communicate with the researchers between sequences.

### 2.3 Experimental procedure

We selected 36 social concepts from the list reported by Anderson (1968), for which we developed short audio definitions controlling for sentence length. We categorized all social concepts following a 2×2 factorial design using the concept dimensions of affect and likableness. First, half of the concepts referred to affective states, making an explicit mention to the emotions of oneself or others (see left panel in Table 1), while the other half involved non-affective mental states, referring to interpersonal behavior that does not explicitly involve any emotional content or state (see right panel in Table1). Second, half of the concepts involved highly likable interpersonal behavior (see upper half in Table 1), whereas the other half described highly unlikable social behavior (see bottom half in Table 1). We kept the number of concepts in each category equivalent, with 9 social concepts in each of the four subcategories (e.g. high affect and low likableness). Each trial began with a fixation period of 250 ms followed by a blank screen for 500 ms (see Figure 1B). Then, participants listened to the definition of a social concept for 3500 ms (e.g. ‘*She gets sad when seeing someone suffering and tries to ease their pain*’; see Table 1 for the complete list of social concept definitions), followed by another period of 2000 ms in which they were instructed to mentally simulate a person of their own choice (e.g. a relative, acquaintance, or famous character) behaving as described in the definition.

**Table 1.**
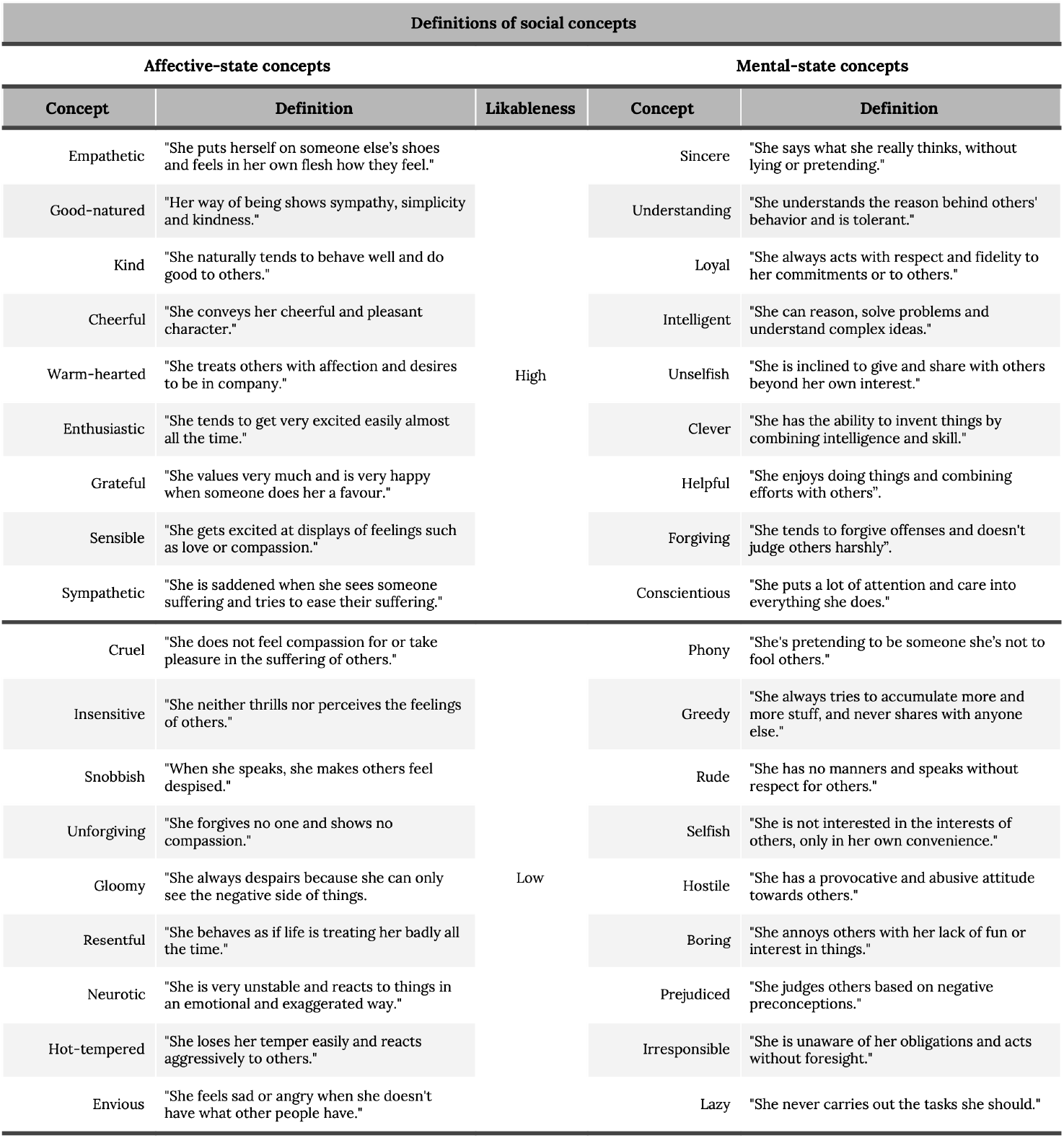
Definitions of social concepts. All concepts used in the experiment followed a 2×2 factorial design. Half of the concepts made an explicit mention to the emotions of oneself or others, while the other half referred to interpersonal behavior that does not explicitly in-volve any emotional content or state. Second, half of the concepts involved highly likable interpersonal behavior, whereas the other half described highly unlikable social behavior.

**Figure 1:**
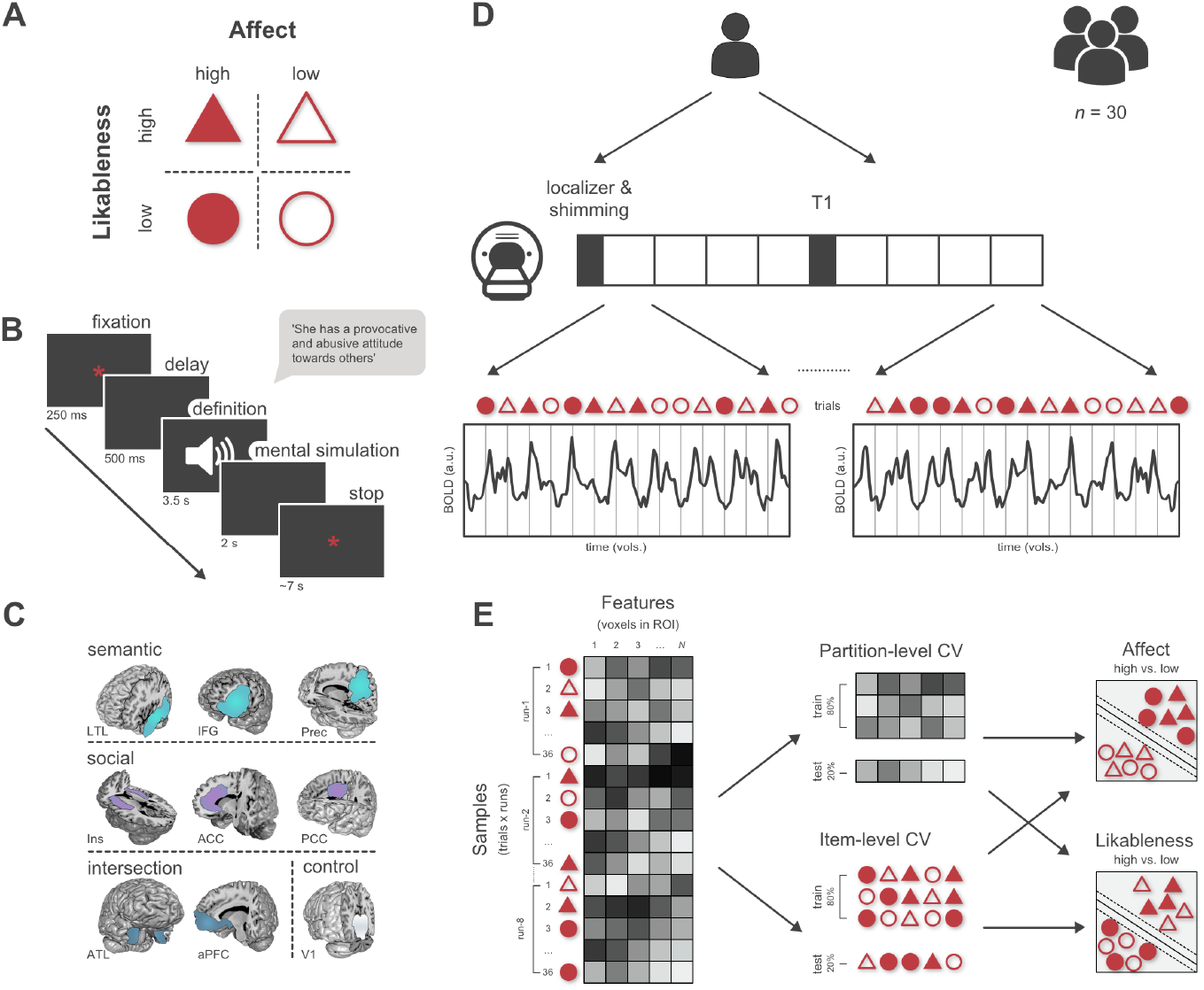
Experiment workflow. (**A**) 36 social concept definitions matched one of our 4 subcategories reflecting a combination of the affect and likableness of the social knowledge. (**B**) Participants listened to the definition of a social concept and were asked to mentally simulate a person behaving the way described in the definition. (**C**) We performed MVPA analyses on the BOLD signals extracted from a set of 9 *a priori* ROIs. (**D**) We acquired one anatomical and 8 functional sequences in a single scanning session. Trial order was randomized between runs. (**E**) After stacking, detrending, and z-scoring the examples, we used two different cross-validation procedures to validate the classification of the affect (high vs. low) and likableness (high vs. low) of social knowledge in separate decoding schemes.

All 36 social concepts were presented in each functional run, with concept order randomized between runs. A run lasted approximately six and a half minutes. To facilitate the estimation of the peak of the HRF across the different trials, we included an additional jitter so that the time between the offset of the current stimulus and the onset of the next audio definition varied between 6 and 8 seconds. The jitter followed a pseudo-exponential distribution resulting in 50% of trials with an ITI of 6 s, 25% of 6.5 s, 12.5% of 7 s and so on. All experimental procedures for stimulus delivery during the mental simulation task were programmed and presented using PsychoPy v.1.83.0.4 (Peirce 2007).

### 2.4 Rating task

Before and after the MRI scanning session, we asked participants to rate the affect and likableness of each concept definition on a scale from 0 to 100. We used these measurements to analyze the test-retest reliability of self-ratings of the concept definitions.

### 2.5 MRI data preprocessing

We first converted all MRI data from DICOM to NIfTI format using MRIConvert (http://lcni.uoregon.edu/downloads/mriconvert). We then preprocessed the MRI data using FEAT 6 (fMRI Expert Analysis Tool) from the FSL suite (FMRIB Software Library; v5.0.9). We removed the first 10 volumes of each functional run to ensure steady-state magnetization. We used FSL’s brain extraction tool 2.1 (BET) to remove non-brain tissue (Smith 2002) and ICA-based Automatic Removal of Motion Artifacts (ICA-AROMA) to identify and remove motion-related artifacts (Pruim et al. 2015). We applied spatial smoothing to the data using a Gaussian kernel of 3 mm FWHM and a high-pass filter with a cutoff of 90 s (estimated using FEAT’s “Estimate High Pass Filter Tool” based on the analysis of the frequency content of the design). All functional images were coaligned to a reference volume from the first run for each participant.

MVPA analyses were conducted in a set of prespecified regions of interest (ROIs) based on previous studies. On the one hand, we derived semantic ROIs from a meta-analysis on semantic processing (Binder, Desai, et al. 2009). These were left-lateralized since extensive evidence shows a strong lateralization of language processing regions, including high-level, multi-modal regions such as the inferior frontal gyrus (Häberling et al. 2016). We also note that a prior meta-analysis (Binder, Desai, et al. 2009) implicated brain regions in the right hemisphere with semantic processing, although semantic-related activity was higher in the left hemisphere. Hence, we focussed on key semantic regions of interest (ROIs) of the left hemisphere in order to constrain the search space of our fMRI analyses.

Accordingly, semantic ROIs included the left lateral temporal lobe (LTL), inferior frontal gyrus (IFG), and precuneus (Prec). On the other hand, we selected a set of social ROIs from a meta-analysis on social information processing (Alcalá-López et al. 2018). These were bilateral since the results from that meta-analysis on the functional connectivity architecture of the social brain showed a strong coupling of the connectivity maps between bilateral regions. Social ROIs included the insula (Ins), and anterior (ACC) and posterior cingulate cortices (PCC). Moreover, we included the anterior temporal lobe (ATL) and anterior prefrontal cortex (aPFC) as ROIs on the intersection of both semantic and social information processing. Finally, we included the primary visual cortex (V1) as a control region. We used FreeSurfer v.6.0.0 for automatic segmentation of the structural images. We then obtained masks for each *a priori* ROI in anatomical space using FreeSurfer’s LookUp Table. After visual inspection of the anatomical masks, we transformed them to each participant’s functional space using FLIRT (Jenkinson, Bannister, et al. 2002; Jenkinson and Smith 2001). All MVPA analyses were performed in native BOLD space.

### 2.6 Multivariate pattern analysis

We conducted multivariate pattern analysis as implemented in the Python libraries scikit-learn (Pedregosa et al. 2011) and PyMVPA (Hanke et al. 2009). We used a linear support vector machine classifier (SVC) to classify the concept category of our definitions in each ROI. We performed two separate binomial classifications, one for the affect (high vs. low affect) and another for the likableness (high vs. low likableness) of social knowledge. These allowed us to characterize which dimension of social knowledge is better represented across the different *a priori* semantic and social ROIs. First, we trained an SVC on examples with randomly shuffled labels and then tested the classifier on the examples labeled appropriately to obtain an empirical estimate of chance level performance. We then compared the average decoding accuracy of each category in the selected ROIs of the canonical semantic network to the selected social-cognitive processing ROIs. Finally, we trained a classifier on examples labeled for the affect condition and tested on examples labeled for the likableness condition, and vice versa. This procedure served as sanity check to ensure that the results from the previous classification analyses reflected information specifically related to the affect and likableness classes, and that an spurious or non-specific signals were driving the effects. Additional details on data preparation, feature selection, classification, cross-validation, and group-level statistics are presented in the following sections.

#### 2.6.1 Data preparation

After preprocessing the MRI data, we used the output generated from PsychoPy during the experimental task to label the relevant scans with an attribute for each binomial classification (i.e. high vs. low affect; high vs. low likableness) for each subject. We then removed invariant features (i.e. voxels whose BOLD activity did not vary throughout the length of a functional run) and stacked the data from all 8 functional runs after z-score normalization and linear detrending (see Figure 1D). Finally, we generated examples for the MVPA analysis by averaging BOLD signal between 5.5 s and 10.5 s after stimulus onset. Given that the audio definitions lasted for about 3.5 seconds, this timeframe was selected to ensure that our BOLD examples for classification contained information from the peak of the HRF associated with processing the content of the definition.

#### 2.6.2 Pattern classification

We used a linear support vector classifier with the default settings provided by scikit-learn (*l2* regularization, *C* = 1.0, *tolerance* = 0.0001) to decode semantic category in each binomial classification. For the cross-validation (CV) of each classification problem, we initially shuffled and resampled the stacked examples multiple times to create 300 partitions of balanced train-test (80%–20%) sets to obtain an unbiased generalization estimate for each ROI (Varoquaux et al. 2017). In this partition-based CV, although training and test sets used independent partitions of scans, the same concept definition could potentially appear in both sets. Hence, we also ran a second CV procedure, in which the training and testing sets used different items (i.e. all scans associated with a selection of concepts were left out for testing). This second item-level CV offers a more robust out-of-sample generalization. To ensure we could compare both CV procedures, we left out two concepts from each subcategory (~20% of all examples) for testing during each iteration (see Figure 1E).

We then used Principal Component Analysis (PCA) with the default settings as provided by scikit-learn to reduce the dimensionality of the data and, thus, the chances of overfitting (Mitchell, Hutchinson, Niculescu, et al. 2004; Pereira et al. 2009). We kept the number of components equal to the number of examples, resulting in all ROIs having the same number of components. Note that PCA was performed on the training set; then the trained PCA was used to extract components in the test set and its classification performance was assessed. This procedure was repeated separately for each of the 300 train-test splitsL. We collected and averaged the corresponding decoding accuracies for each participant in each ROI.

#### 2.6.3 Statistics

We first assessed the test-retest reliability of the participants’ subjective ratings of our concept definitions before and after the scanning session. We calculated the intraclass correlation coefficient (ICC) and its 95% confidence intervals as implemented in Python in the Pingouin library (Vallat 2018).

We then performed group-level statistics on the MVPA classification accuracies across subjects for each ROIs using the open-source statistical program JASP (JASP Team, 2019, jasp-stats.org). We used paired t-tests to compare the decoding accuracy at each ROI to the estimated chance level performance. Then, we used two repeated-measures ANOVAs with one factor (ROI) to determine statistically significant differences in the decoding accuracy of each ROI for the classification of affect and likableness, respectively. We then used a two-way repeated-measures ANOVA with two factors (ROI * concept category) to determine whether one of our social knowledge dimensions (affect vs. likableness) could be decoded with higher classification accuracy than the other. Finally, we performed a paired t-test between the average decoding accuracy in canonical semantic regions (i.e. LTL, IFG, and Prec) and the average accuracy in regions from the social brain (i.e. Ins, ACC, PCC) across subjects.

All analyses were performed on the resulting decoding accuracies using both the partition-level and item-level cross-validation procedures. All code used for data processing and analysis is available on GitHub (https://github.com/danalclop/socialcon).

## 3 Results

### 3.1 Behavioral results

Subjective ratings of the concept definitions showed that participants categorized social concepts in terms of their affect and likableness as expected based on their normative definitions (Anderson 1968; see Figure 2). A paired *t*-test confirmed that ratings of affect among the concepts defining affective states (*M* = 68.514, *SD* = 12.614) were significantly higher than those describing non-affective mental states (*M* = 45.813, *SD* = 13.442; *t*_(29)_ = 8.026, *p* < 0.001, *d* = 1.465, *logBF10* = 14.35). Similarly, ratings of likableness among the concepts defined as socially desirable (*M* = 84.472, *SD* = 6.580) were significantly higher than those selected as highly unlikable (*M* = 15.413, *SD* = 8.285; *t*_(29)_ = 30.382, *p* < 0.001, *d* = 5.547, *logBF10* = 46.69). Such difference between ratings of high vs. low affect (*M* = 22.701, *SD* = 15.493) was smaller than the difference between ratings of high vs. low likableness (*M* = 69.060, *SD* = 12.450; *t*_(539)_ = −13.925, *p* < 0.001, *d* = −2.542, *logBF10* = 26.42). Note that a *logBF10* greater than 2.2 is considered an overwhelming support in favour of the alternative hypothesis (Kass and Raftery 1995). This suggests that the social desirability of others’ behavior is more salient for the representation of social knowledge than affect.

**Figure 2:**
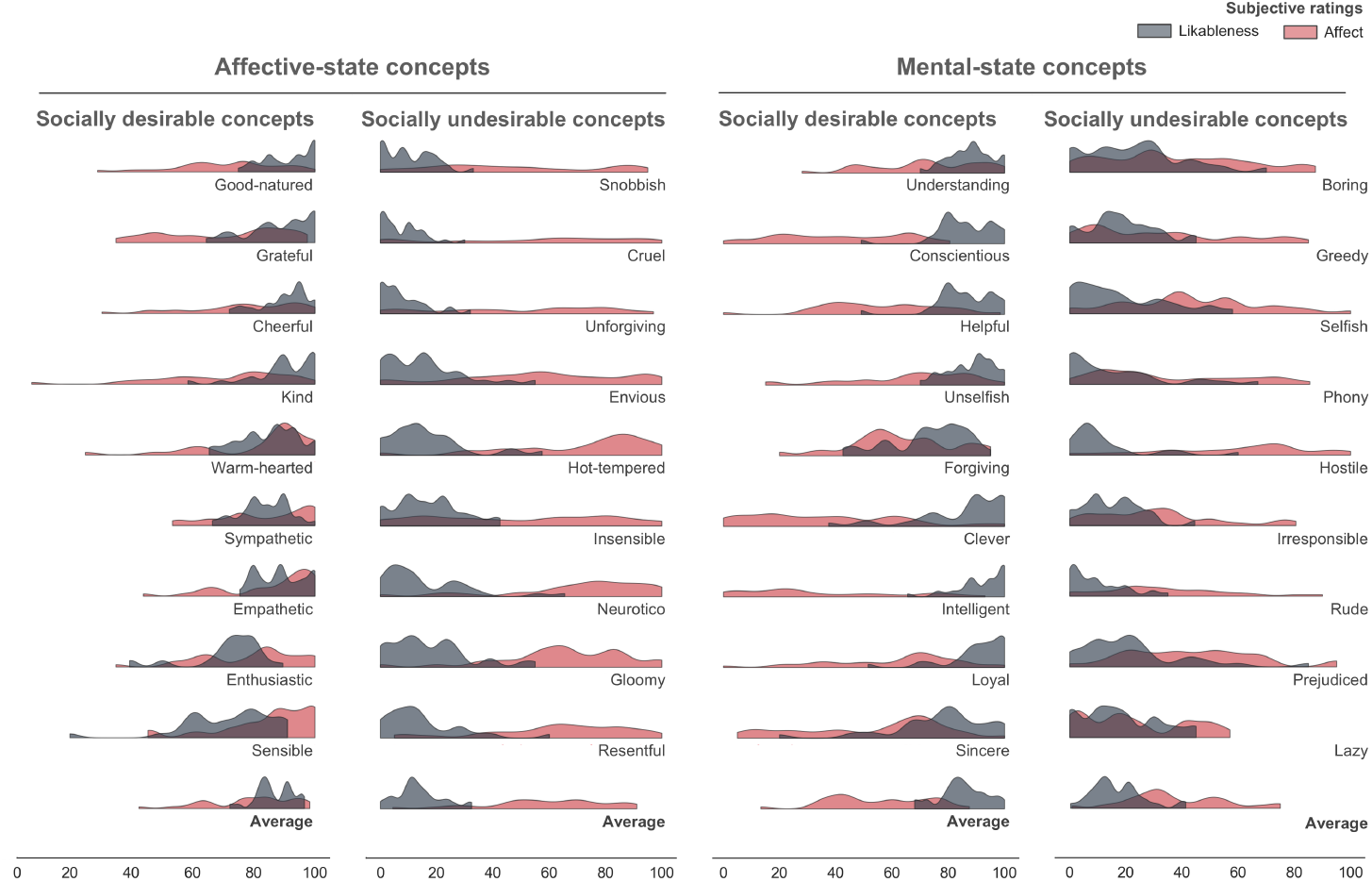
Distributions of ratings of social concepts. Participants read each concept definition and rated the extent to which the described behavior involved the emotions of oneself or others (affect; *red*) as well as whether such behavior was socially desirable (likableness; *gray*) on a scale from 0 (very non-affective; very unlikable) to 100 (very affective; very likable).

This is congruent with the results from the test-retest repeatability analysis. The intraclass correlation coefficient showed that the reliability of the ratings before and after the scanning session was fair for affect (ICC = 0,47; 95% CI [0,36-0,60]) and excellent for likableness (ICC = 0,93; 95% CI [0,89-0,96]).

### 3.2 Brain imaging results

As mentioned above, we used two cross-validation procedures for the binomial classification analyses. In the partition-level CV, we used independent partitions of the scans for training and testing (i.e. training occurred within partitions containing 80% of the data; then, we tested in the remaining 20%). Nonetheless, scans in both sets could refer to the same concepts. In the second cross-validation procedure, the training and testing partitions did not include scans corresponding to the same concepts, hence providing a better estimate of out-of-sample generalization.

A set of paired *t*-tests confirmed that the decoding accuracy of both affect and likableness was significantly above the empirical chance level in all ROIs when using partition-level CV (see Figure 3A). We then repeated the decoding analysis with the second, item-level CV procedure that allowed testing the generalizability of the brain representations of social concepts. While decoding accuracy of affect was significantly higher than chance in all ROIs when using item-level CV, it did not differ significantly from chance in the ATL (*t*_(29)_ = 2.248, *p* = 0.037, *d* = 0.410, *logBF10* = 0.529), Ins (*t*_(29)_ = 2.244, *p* = 0.033, *d* = 0.410, *logBF10* = 0.522), and V1 (*t*_(29)_ = −1.981, *p* = 0.057, *d* = −0.362, *logBF10* = 0.073) for the classification of likableness. Correct classification rates using partition-level CV, chance performance, and summary statistics for each ROI are presented in Table 2 (see Table 4 in the supplementary materials for results using item-level CV).

**Figure 3:**
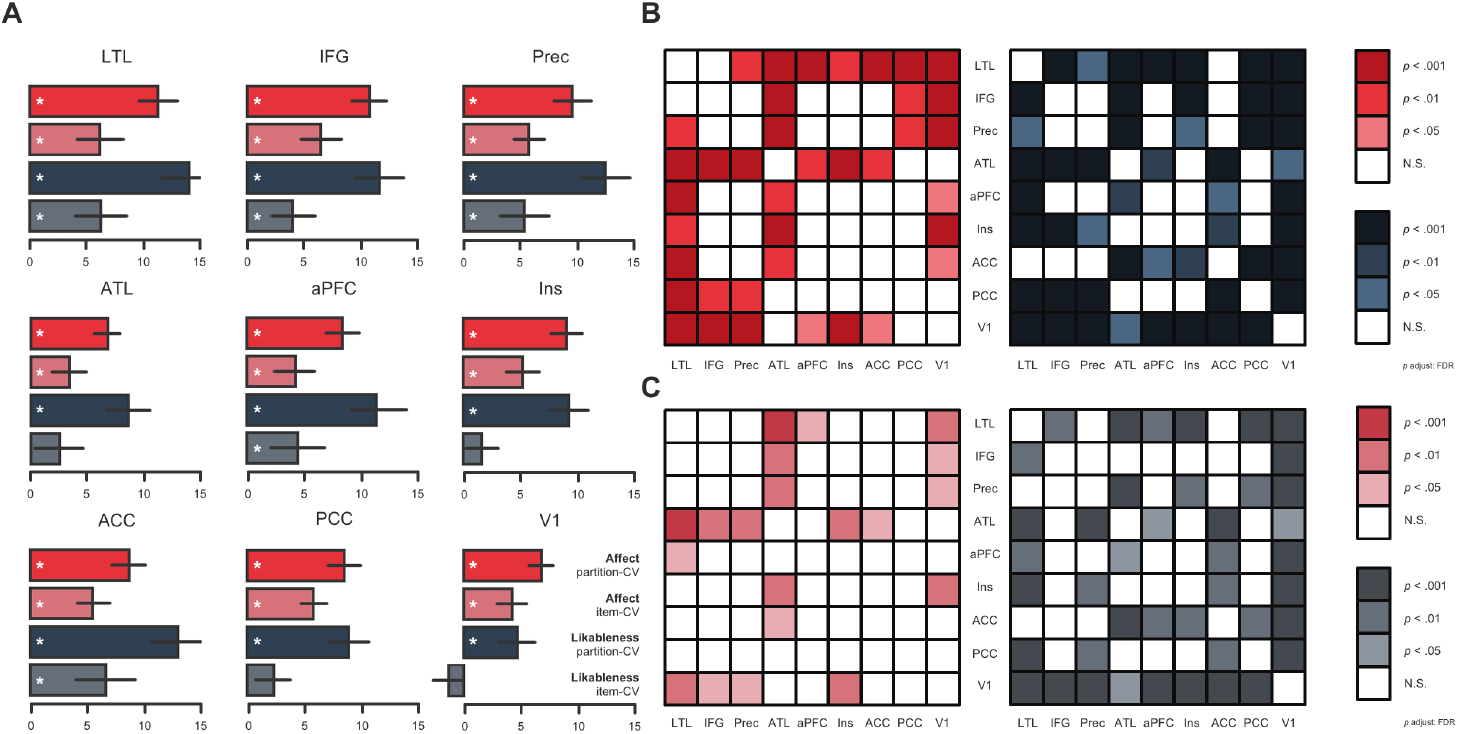
Decoding accuracies of social concepts across the brain. (**A**) Classification accuracy in all ROIs was significantly above chance (white asterisks mark significant differences using a threshold of *α* = 0.001 after correction for multiple comparisons) when using a partition-level CV, whereas the classifier performed at chance when decoding likableness in the ATL, Ins, PCC, and V1 using an item-level CV. The origin value in the horizontal axis marks the empirically-estimated chance level for each classification problem (partition-level CV: *M* = 53.40%; item-level CV: *M* = 53.61%). (**B**) *Post hoc* paired *t*-tests of two separate repeated-measures ANOVAs with one factor (ROI) showing significant differences between ROIs in the classification of affect (*left*) and likableness (*right*) of social knowledge when using a partition-level CV and (**C**) when using an item-level CV procedure. Note that color saturation indicates cross-validation type and color hue marks distinct thresholds of statistical significance in panels (B) and (C).

**Table 2.** Binomial classification of the affect and likableness of social knowledge. Correct classification rates, chance performance, and summary statistics for the binomial classification problems of affect (high vs. low; *top*) and likableness (high vs. low; *bottom*) using a partition-level cross-validation.

| Affect |  |  |  | 95% CI of mean difference |  |  |  |  |  |
| --- | --- | --- | --- | --- | --- | --- | --- | --- | --- |
| ROI | Estimate | Chance | Mean difference | Lower | Upper | SE difference | t | P<sub>FDR</sub> | Cohen's d |
| LTL | 0.653 | 0.541 | 0.112 | 0.094 | 0.131 | 0.009 | 12.469 | < .001 | 2.277 |
| IFG | 0.637 | 0.530 | 0.107 | 0.092 | 0.122 | 0.007 | 14.422 | < .001 | 2.633 |
| Prec | 0.634 | 0.539 | 0.095 | 0.079 | 0.112 | 0.008 | 11.682 | < .001 | 2.133 |
| ATL | 0.594 | 0.526 | 0.068 | 0.056 | 0.080 | 0.006 | 11.451 | < .001 | 2.091 |
| aPFC | 0.621 | 0.538 | 0.083 | 0.068 | 0.099 | 0.007 | 11.269 | < .001 | 2.057 |
| Ins | 0.628 | 0.538 | 0.090 | 0.076 | 0.105 | 0.007 | 13.172 | < .001 | 2.405 |
| ACC | 0.623 | 0.536 | 0.086 | 0.071 | 0.102 | 0.008 | 11.393 | < .001 | 2.080 |
| PCC | 0.612 | 0.527 | 0.085 | 0.070 | 0.100 | 0.007 | 11.570 | < .001 | 2.112 |
| VI | 0.598 | 0.530 | 0.068 | 0.057 | 0.080 | 0.006 | 12.301 | < .001 | 2.246 |

TABLE 2
| Likableness |  |  |  | 95% CI of mean difference |  |  |  |  |  |
| --- | --- | --- | --- | --- | --- | --- | --- | --- | --- |
| ROI | Estimate | Chance | Mean difference | Lower | Upper | SE | t | P<sub>FDR</sub> | Cohen's d |
| LTL | 0.682 | 0.543 | 0.139 | 0.113 | 0.166 | 0.013 | 10.634 | < .001 | 1.942 |
| IFG | 0.651 | 0.535 | 0.116 | 0.094 | 0.138 | 0.011 | 10.799 | < .001 | 1.972 |
| Prec | 0.659 | 0.535 | 0.124 | 0.101 | 0.147 | 0.011 | 10.952 | < .001 | 2.000 |
| ATL | 0.608 | 0.523 | 0.085 | 0.064 | 0.106 | 0.010 | 8.290 | < .001 | 1.514 |
| aPFC | 0.645 | 0.532 | 0.113 | 0.087 | 0.140 | 0.013 | 8.833 | < .001 | 1.613 |
| Ins | 0.631 | 0.539 | 0.092 | 0.075 | 0.109 | 0.008 | 11.051 | < .001 | 2.018 |
| ACC | 0.666 | 0.537 | 0.129 | 0.104 | 0.154 | 0.012 | 10.648 | < .001 | 1.944 |
| PCC | 0.622 | 0.533 | 0.089 | 0.070 | 0.108 | 0.009 | 9.633 | < .001 | 1.759 |
| VI | 0.580 | 0.533 | 0.047 | 0.031 | 0.063 | 0.008 | 5.910 | < .001 | 1.079 |

We then used two repeated-measures ANOVAs with ROI as a factor to analyze significant differences in decoding accuracy among ROIs for each classification problem.

On the one hand, using the partition-level CV we found a main effect of ROI for both the classification of affect (*F*_(8, 232)_ = 19.505, *p* < 0.001, *ω*^2^ = 0.244, *logBF10* = 44.654, *R*^2^ = 0.547) and likableness (*F*_(8, 232)_ = 30.888, *p* < 0.001, *ω*^2^ = 0.267, *logBF10* = 67.704, *R*^2^ = 0.686). *Post hoc* tests showed that the decoding accuracy in the LTL was significantly higher than the rest of the ROIs except for the IFG for the classification of affect (*t*_(29)_ = 2.417, *p* = 0.266, *d* = 0.441, *logBF10* = 0.840), and except for the ACC for the classification of likableness (*t*_(29)_ = 2.038, *p* = 0.424, *d* = 0.366, *logBF10* = 0.167; see Figure 3B). Although all ROIs classified the likableness of social concepts better than the control ROI V1, the ATL did not classify the affect of concepts significantly better than V1 (*t*_(29)_ = 0.758, *p* = 1.00, *d* = 0.138, *logBF10* = 1.374).

On the other hand, using the item-level CV we also found a main effect of ROI for the classification of both affect (*F*_(8, 232)_ = 8.195, *p* < 0.001, *ω*^2^ = 0.240, *logBF10* = 16.386, *R*^2^ = 0.422) and likableness (*F*_(8, 232)_ = 23.531, *p* < 0.001, *ω*^2^ = 0.218, *logBF10* = 53.163, *R*^2^ = 0.660). However, *post hoc* tests on the classification of affect pointed that decoding accuracy in the LTL was only significantly higher than the ATL (*t*_(29)_ = 4.896, *p* = 0.001, *d* = 0.894, *logBF10* = 6.540), aPFC (*t*_(29)_ = 3.949, *p* = 0.014, *d* = 0.721, *logBF10* = 4.190), and V1 (*t*_(29)_ = 4.314, *p* = 0.005, *d* = 0.788, *logBF10* = 5.082; see Figure 3C). The control ROI V1 also showed a significantly lower accuracy than the IFG (*t*_(29)_ = 3.546, *p* = 0.036, *d* = 0.647, *logBF10* = 3.236), Prec (*t*_(29)_ = 3.662, *p* = 0.029, *d* = 0.669, *logBF10* = 3.506), and Ins (*t*_(29)_ = 4.586, *p* = 0.003, *d* = 0.837, *logBF10* = 5.760; see Table 5 & Table 6 in the supplementary materials for all *post hoc* tests of the classification of affect when using partition-level and item-level CV, respectively). For the classification of the likableness of social concepts, the LTL decoded significantly better than the rest of the ROIs except the Prec (*t*_(29)_ = 2.877, *p* = 0.102, *d* = 0.525, *logBF10* = 1.754) and ACC (*t*_(29)_ = 0.349, *p* = 1.00, *d* = 0.064, *logBF10* = −1.581; see Figure 3C), and all ROIs outperformed the control ROI V1 (see Table 5 & Table 7 in the supplementary materials for all *post hoc* tests of the classification of likableness when using partition-level and item-level CV, respectively).

In sum, we found that our selection of semantic and social regions contains detailed information on the meaning of social concepts that allows a linear classifier to distinguish between subcategories based on their affect and likableness. These results were supported by a second, more stringent cross-validation procedure, which yielded results almost identical to the typical 5-fold CV procedure using stratified random partitions of all brain scans (see Figure 5 in the supplementary materials for a comparison of the distribution of all decoding accuracies across concept dimensions and cross-validation procedures).

To further evaluate whether our ROIs showed a bias towards the decoding of the affect or likableness of social knowledge, we performed a repeated-measures ANOVA with two factors: ROI and concept dimension (i.e. affect vs. likableness). We found a main effect of dimension for the partition-level cross-validation (*F*_(1, 29_) = 7.801, *p* = 0.009, *ω*^2^ = 0.054, *logBF10* = 20.064, *R*^2^ = 0.302), showing that the decoding accuracy of the likableness dimension was higher than the affect dimension (see Table 3). Moreover, we found an interaction between ROI and dimension (*F*_(1, 29)_ = 7.640, *p* < .001, *ω*^2^ = 0.056, *logBF10* = 8.558), indicating that decoding of affect and likableness information varied across ROIs (see Fig. 5 in the supplementary materials for the complete distribution of classification accuracies). *Post hoc* paired *t*-tests between the decoding accuracies of affect and likableness showed that this effect was driven by the ACC, which showed a preference for the likableness dimension (decoding accuracy of likableness: *M* = 0.666, *SD* = 0.058; affect: *M* = 0.623, *SD* = 0.034; *t*_(29)_ = 4.461, *p* = 0.001, *d* = 0.814, *logBF10* = 5.448; see Table 8 for decoding accuracy and summary statistics of all ROIs using partition-level cross-validation). Conversely, when using the item-level cross-validation, the main effect of dimension was not statistically significant (*F*_(1, 29)_ = 2.938, *p* = 0.097, *ω*^2^ = 0.043, *logBF10* = 13.351, *R*^2^ = 0.183), whereas the interaction effect remained significant (*F*_(1, 29)_ = 7.281, *p* < .001, *ω*^2^ = 0.105, *logBF10* = 6.436). In this case, *post hoc* paired *t*-tests showed that the effect was driven by the insula but in the opposite direction: the classification of affect outperformed that of likableness (decoding accuracy of likableness: *M* = 0.559, *SD* = 0.032; affect: *M* = 0.595, *SD* = 0.023; *t*_(29)_ = −4.623, *p* <.001, *d* = 0.844, *logBF10* = 5.854). Interestingly, we found a similar result in the control ROI V1 (decoding accuracy of likableness: *M* = 0.521, *SD* = 0.029; affect: *M* = 0.571, *SD* = 0.025; *t*_(29)_ = −6.310, *p* <.001, *d* = −1.152, *logBF10* = 10.132; see Table 8 in the supplementary materials for decoding accuracy and summary statistics of all ROIs using item-level cross-validation). Nevertheless, both regions failed to classify the likableness of social knowledge significantly above chance when using this CV procedure.

**Table 3.** Comparison between the classification accuracies of the affect and likableness of social knowledge. Correct classification rates and summary statistics for the contrast between the classification of likableness (high vs. low) vs. affect (high vs. likableness) using a partition-level cross-validation.

| ROI | Likableness | Affect | Mean difference | 95% CI of mean difference | | SE | t | $p_{FDR}$ | Cohen's $d$ |
| --- | --- | --- | --- | --- | --- | --- | --- | --- | --- |
|  |  |  |  | Lower | Upper |  |  |  |  |
| LTL | 0.682 | 0.653 | 0.029 | 0.009 | 0.049 | 0.010 | 2.931 | 0.016 | 0.535 |
| IFG | 0.651 | 0.637 | 0.013 | -0.006 | 0.032 | 0.009 | 1.429 | 0.210 | 0.261 |
| Prec | 0.659 | 0.634 | 0.024 | 0.010 | 0.039 | 0.007 | 3.420 | 0.009 | 0.624 |
| ATL | 0.608 | 0.594 | 0.014 | -0.003 | 0.032 | 0.009 | 1.671 | 0.158 | 0.305 |
| aPFC | 0.645 | 0.621 | 0.024 | 0.004 | 0.044 | 0.010 | 2.500 | 0.032 | 0.456 |
| Ins | 0.631 | 0.628 | 0.003 | -0.008 | 0.014 | 0.005 | 0.547 | 0.589 | 0.100 |
| ACC | 0.666 | 0.623 | 0.044 | 0.024 | 0.064 | 0.010 | 4.461 | 0.001 | 0.814 |
| PCC | 0.622 | 0.612 | 0.010 | -0.005 | 0.025 | 0.007 | 1.352 | 0.211 | 0.247 |
| V1 | 0.580 | 0.598 | -0.018 | -0.031 | -0.006 | 0.006 | -3.095 | 0.012 | -0.565 |

Next, we asked whether social knowledge was differentially represented in the a priori ROIs defined as canonical semantic (LTL, IFG, and Prec) compared with those defined as social (Ins, ACC, and PCC) according to previous meta-analyses (Alcalá-López et al. 2018; Binder, Desai, et al. 2009). To this end, we averaged the decoding accuracy within semantic and within social ROIs and conducted paired *t*-tests (see Figure 4). Average classification accuracy was significantly higher in semantic ROIs compared to social ROIs for both affect (*M* = 0.642±0.029 in semantic ROIs vs. *M* = 0.621±0.022 in social ROIs; *t*_(29)_ = 5.590, *p* < .001, *d* = 1.021, *logBF10* = 8.305) and likableness (*M* = 0.664±0.050 in semantic ROIs vs. *M* = 0.640±0.034 in social ROIs; *t*_(29)_ = 5.113, *p* < .001, *d* = 0.933, *logBF10* = 7.090) using partition-level cross-validation. These results were supported by the item-level cross-validation procedure: decoding of concept class was higher in semantic ROIs compared to social ROIs for both affect (*M* = 0.599±0.026 in semantic ROIs vs. *M* = 0.591±0.022 in social ROIs; *t*_(29)_ = 2.519, *p* = 0.018, *d* = 0.460, *logBF10* = 1.033) and likableness (*M* = 0.591±0.047 in semantic ROIs vs. *M* = 0.574±0.035 in social ROIs; *t*_(29)_ = 4.133, *p* < .001, *d* = 0.755, *logBF10* = 4.639).

**Figure 4:**
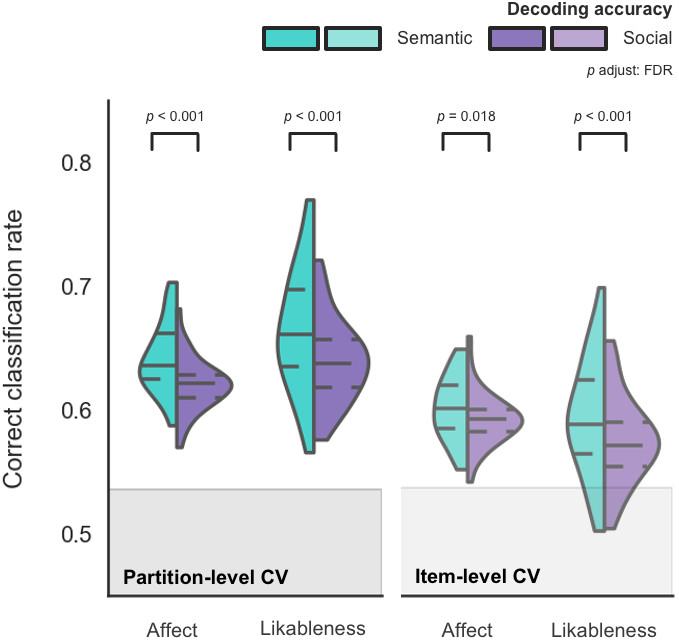
Decoding accuracy of social concepts in semantic vs. social ROIs. Average classification accuracy in semantic ROIs was significantly higher than in social ROIs using both partition- and item-level cross-validation procedures. The shaded area indicates the average empirically-estimated chance level (*M* = 0.53).

As a final verification check, we addressed whether the neural pattern underlying each concept dimension of social knowledge was distinct. Accordingly, we tested whether an SVC trained to discriminate the affect dimension (high vs. low) generalized to classify examples in the likableness condition, and vice versa, using the item-level CV. This analysis allowed us to rule out the possibility that classification accuracy of both social dimensions was mediated by a common factor. Results showed that training in affect and testing in likableness produced decoding accuracies at chance level (*M* = 0.5013) and, reciprocally, training in likableness and testing in affect also resulted in chance level classification (*M* = 0.501; see Table 9).

## 4 Discussion

The present fMRI study investigated how social knowledge is represented in the brain. In particular, we asked whether the affect and likableness of interpersonal behavior are relevant features that the brain uses to construct such representations. We used multi-variate classification analyses from BOLD signals in semantic (LTL, IFG, and Prec) and social-cognitive processing regions (Ins, ACC, and PCC) to understand how these key dimensions of social concepts are represented in the human brain. The main goal of this study was to test whether activity patterns in functionally distinct brain regions contain information that is detailed enough to allow a linear classifier to distinguish between sub-categories of social concepts defined *a priori*.

We found that most of these regions can decode above chance the affect and lik-ableness dimensions of social knowledge, with slight differences depending on the cross-validation procedure used to validate the performance of the classifier. Decoding accuracy was particularly high in the lateral temporal lobe (LTL) for the classification of both the affect and likableness of interpersonal behavior. On the contrary, the visual cortex, which we included in the analysis as a control region, proved to be the least accurate of the ROIs measured when decoding social knowledge. Nevertheless, even in the visual cortex the classification accuracy was significantly above chance–except for the classification of lik-ableness using an item-level CV. Interestingly, this result is in line with a recent study that used a convolutional neural network to decode different emotion categories from visual stimuli (Kragel et al. 2019). The results from this study also showed that it is possible to use activity patterns in the visual cortex to discriminate between emotion categories as well as the valence of the images. Although we used auditory stimuli, it is likely that visual representations were co-activated since participants were instructed to engage in mental simulation of the auditory definitions. It is known that mental imagery engages the early visual cortex (Pearson et al. 2015). Hence, our results also support the idea that early sensory areas participate in the representation of abstract concepts, at least to a certain degree. These results are in keeping with the view that the representation of concrete and abstract concepts relies on partially overlapping mechanisms (Borghi et al. 2019) and that both may rely to some extent in domain-specific processing.

We also evaluated whether the concept dimensions of affect and likableness played different roles in the brain representation of social knowledge. We found that the ACC decoded the likableness of interpersonal behavior significantly better than its affect when using a partition-level CV. This is congruent with the self-reported ratings we obtained from the participants, which indicate that ratings of likableness were more concentrated in the extreme values of the distribution for corresponding high and low normative values (Anderson 1968). Moreover, these ratings of likableness were consistent with recent replication studies (Chandler 2018; Dumas et al. 2002). Nevertheless, activity patterns in the insula allowed for decoding the affect of social concepts significantly better when using the more stringent cross-validation procedure involving out-of-sample generalization across new concepts. One possibility is that the different cross-validation procedures may tap onto different levels of abstraction of the representations supported by the ACC and the Ins. Note that both the ACC and Ins were included as regions of interest precisely due to their fundamental role in the processing of socially-relevant information. For instance, during social evaluation tasks, the ACC has been implicated in the detection of positively valenced attributes that relate to the self (Sharot et al. 2007) and also others (Hughes and Beer 2012), while there may exist specialised processes in the ACC for the detection of valence and the subjective reporting of self-rewarding attributes during social evaluation (Rigney et al. 2017). Additional evidence further indicates suggests that ACC and Ins are implicated in processing salient cues related to the self (Perini et al. 2018). Our results suggest that both the ACC and Ins, as part of the salience network (Uddin 2015), may be involved in representing these distinct aspects of social knowledge related to the likableness and affect dimensions. Notably, this was observed in a context that did not require participants to perform overt responses to items in a social experimental task setting, rather we only required mental simulation of social situations based on personal, idiosyncratic experiences (Soto et al. 2020). These findings need further exploration in future studies aimed specifically to better understand the role of the ACC and Ins in the simulation of social contexts.

A further goal of this study was to assess the contribution of canonical semantic regions and social cognitive regions in the brain representation of social concepts. Prior research has lent support for the view that both semantic and social systems play a role in the brain representation of concept knowledge. Brain activity evoked by abstract compared to concrete concepts during different semantic, lexical, and sentence comprehension tasks consistently shows increased activation in a left-lateralized semantic network including inferior frontal and anterior and lateral temporal areas (J. Wang et al. 2010). On the other hand, a recent fMRI study found that distributed activity patterns in the superior and inferior parietal lobe, as well as in the inferior and middle frontal gyri, can be used to decode the intentions behind a set of reach-to-grasp movements performed by another individual (Koul et al. 2018). These brain regions are commonly referred to as the putative *mirror neuron system*, which led the researchers to conclude that the same regions that engage in the perception and action of specific goal-directed behaviors are also responsible for the representation of the intentions of others. Although we found that both sets of ROIs decoded social knowledge above chance level, our results indicate that average decoding accuracy in semantic regions, including the LTL, IFG, and Prec, was significantly higher than in social-cognitive processing areas such as the Ins, ACC, and PCC. Moreover, this finding was robust over both cross-validation procedures. Further, we implemented a feature selection procedure based on PCA that ensured that the dimensionality of the data fed to the classifier was similar across ROIs. Together with our finding that modality-specific systems in the visual cortex contain relevant information to decode social concepts, these results support the hypothesis that sensorimotor features are important for the representation of both concrete and abstract concepts, but critically the latter further rely on language and social-cognitive processes, possibly due to their larger variability and semantic difficulty (Borghi et al. 2019).

It is worth noting that although the ATL has received much attention in recent years due to its involvement in the processing of abstract concepts (Binney et al. 2016; Hoffman et al. 2015; Y. Wang et al. 2017), our results do not place this brain region in a privileged position when it comes to the representation of the likableness and the affective dimension of social information. We show that a linear classifier can indeed use the distributed activity patterns within the ATL to decode each of these dimensions (see Figure 3), however, decoding from other semantic and social ROIs outperformed the ATL in these classification problems. Nevertheless, we are mindful that the MRI signal tends to be weak in this area and hence the ATL results must be taken with caution; we cannot rule out the higher classification accuracies in other regions compared to the ATL may be to signal loss in this area. In a similar fashion, the MPFC was previously implicated in the representation of trait knowledge about other individuals (Ma et al. 2014). In contrast, we find that the aPFC does not show a preferential role in the representation of social knowledge. Activation patterns in this region allowed the linear classifier to significantly decode the affect and likableness of social concepts, disregarding the type of cross-validation procedure used. However, as shown in Figure 3, the lateral temporal lobe showed higher classification accuracy than the aPFC.

Recent studies have suggested a variety of models to explain the conceptual dimensions underlying the representation of social knowledge. Tamir and colleagues used fMRI and representational similarity analyses to study the brain representation of internal states of other individuals (Tamir et al. 2016). Participants had to consider up to 60 different internal states (e.g. *drunkenness* or *satisfaction*). Then, they used representational similarity analyses to demonstrate how a theoretical model of how different social dimensions inter-relate can explain the brain response when subjects consider the internal states of others. Their results showed that three concept dimensions, namely rationality, social impact, and valence, significantly explained a great deal of the variance in the brain representations of other people’s mental states (Tamir et al. 2016). Here we showed how the affect and likableness dimensions of social knowledge are represented in multi-voxel patterns across putative language semantic and social processing regions. Concerning the differences in the number of concept dimensions in Tamir and cols. (2016) and our study, one possibility is that the dimension of likableness or social desirability that we identified here results from the combination of the ‘sociality’ component of the social impact dimension identified by Tamir and cols. (2016).

In sum, the results from our study are not congruent with a modular view of the representation of abstract concept knowledge about other individuals. Rather, they underscore the distributed nature of the brain representation of social concepts, which, although may be partially grounded on sensorimotor features and social cognitive systems, it relies heavily on processing in language-semantic systems.

## Acknowledgements

We acknowledge support from the Basque Government through the BERC 2018-2021 program, from the Spanish Ministry of Economy and Competitiveness, through the ‘Severo Ochoa’ Programme for Centres/Units of Excellence in R & D (SEV-2015-490).

## A Supplementary materials

**Table 4.**
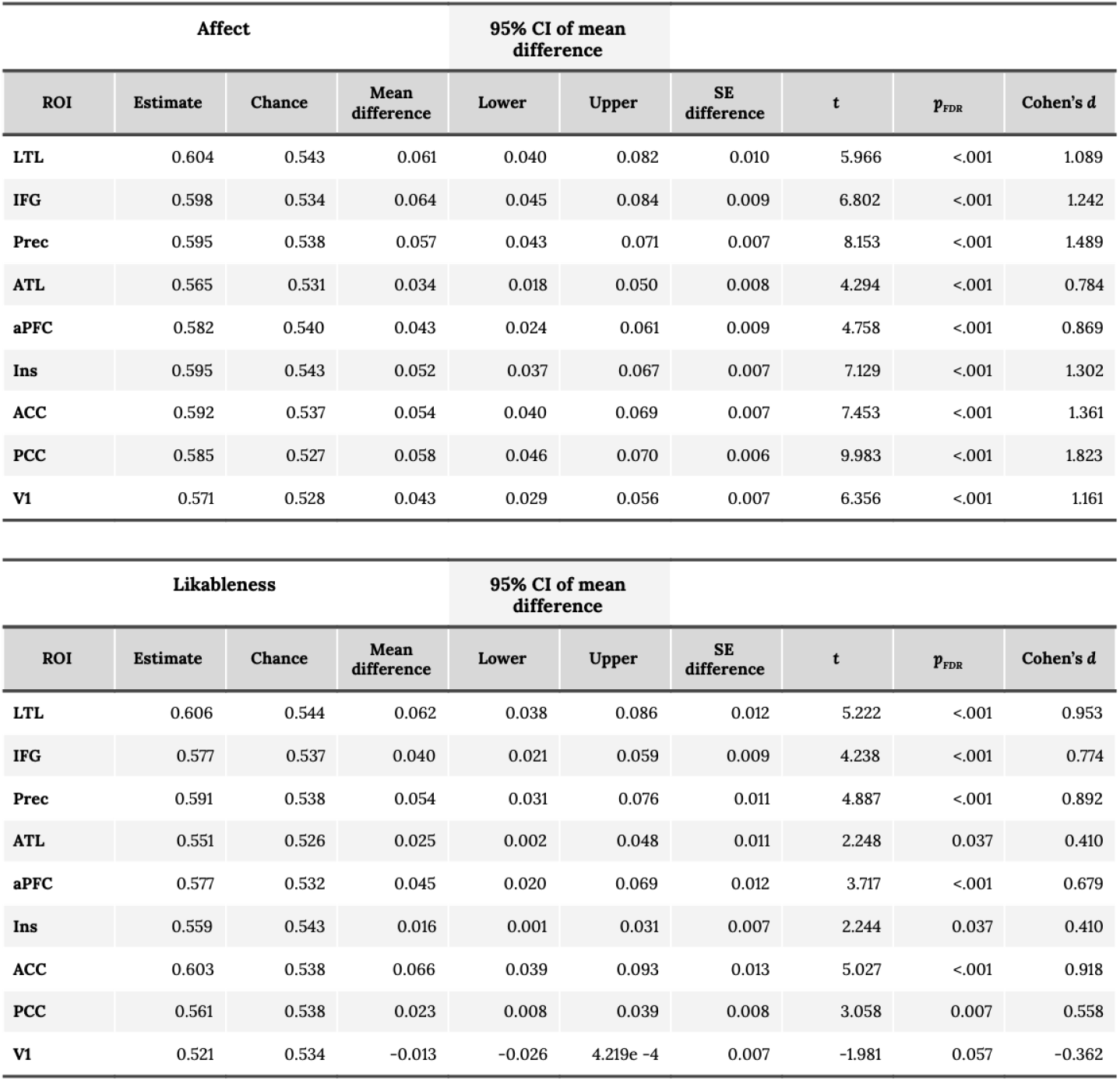
Binomial classification of the affect and likableness of social knowledge. Correct classification rates, chance performance, and summary statistics for the binomial classification problems of affect (high vs. low; *top*) and likableness (high vs. low; *bottom*) using an item-level cross-validation.

| Affect |  |  |  | 95% CI of mean difference |  |  |  |  |  |
| --- | --- | --- | --- | --- | --- | --- | --- | --- | --- |
| ROI | Estimate | Chance | Mean difference | Lower | Upper | SE difference | t | p<sub>FDR</sub> | Cohen's d |
| LTL | 0.604 | 0.543 | 0.061 | 0.040 | 0.082 | 0.010 | 5.966 | <.001 | 1.089 |
| IFG | 0.598 | 0.534 | 0.064 | 0.045 | 0.084 | 0.009 | 6.802 | <.001 | 1.242 |
| Prec | 0.595 | 0.538 | 0.057 | 0.043 | 0.071 | 0.007 | 8.153 | <.001 | 1.489 |
| ATL | 0.565 | 0.531 | 0.034 | 0.018 | 0.050 | 0.008 | 4.294 | <.001 | 0.784 |
| aPFC | 0.582 | 0.540 | 0.043 | 0.024 | 0.061 | 0.009 | 4.758 | <.001 | 0.869 |
| Ins | 0.595 | 0.543 | 0.052 | 0.037 | 0.067 | 0.007 | 7.129 | <.001 | 1.302 |
| ACC | 0.592 | 0.537 | 0.054 | 0.040 | 0.069 | 0.007 | 7.453 | <.001 | 1.361 |
| PCC | 0.585 | 0.527 | 0.058 | 0.046 | 0.070 | 0.006 | 9.983 | <.001 | 1.823 |
| V1 | 0.571 | 0.528 | 0.043 | 0.029 | 0.056 | 0.007 | 6.356 | <.001 | 1.161 |

TABLE 4 **Binomial classification of the affect and likableness of social knowledge.**
| Likableness |  |  |  | 95% CI of mean difference |  |  |  |  |  |
| --- | --- | --- | --- | --- | --- | --- | --- | --- | --- |
| ROI | Estimate | Chance | Mean difference | Lower | Upper | SE difference | t | p<sub>FDR</sub> | Cohen's d |
| LTL | 0.606 | 0.544 | 0.062 | 0.038 | 0.086 | 0.012 | 5.222 | <.001 | 0.953 |
| IFG | 0.577 | 0.537 | 0.040 | 0.021 | 0.059 | 0.009 | 4.238 | <.001 | 0.774 |
| Prec | 0.591 | 0.538 | 0.054 | 0.031 | 0.076 | 0.011 | 4.887 | <.001 | 0.892 |
| ATL | 0.551 | 0.526 | 0.025 | 0.002 | 0.048 | 0.011 | 2.248 | 0.037 | 0.410 |
| aPFC | 0.577 | 0.532 | 0.045 | 0.020 | 0.069 | 0.012 | 3.717 | <.001 | 0.679 |
| Ins | 0.559 | 0.543 | 0.016 | 0.001 | 0.031 | 0.007 | 2.244 | 0.037 | 0.410 |
| ACC | 0.603 | 0.538 | 0.066 | 0.039 | 0.093 | 0.013 | 5.027 | <.001 | 0.918 |
| PCC | 0.561 | 0.538 | 0.023 | 0.008 | 0.039 | 0.008 | 3.058 | 0.007 | 0.558 |
| V1 | 0.521 | 0.534 | -0.013 | -0.026 | 4.219e -4 | 0.007 | -1.981 | 0.057 | -0.362 |

**Figure 5:**
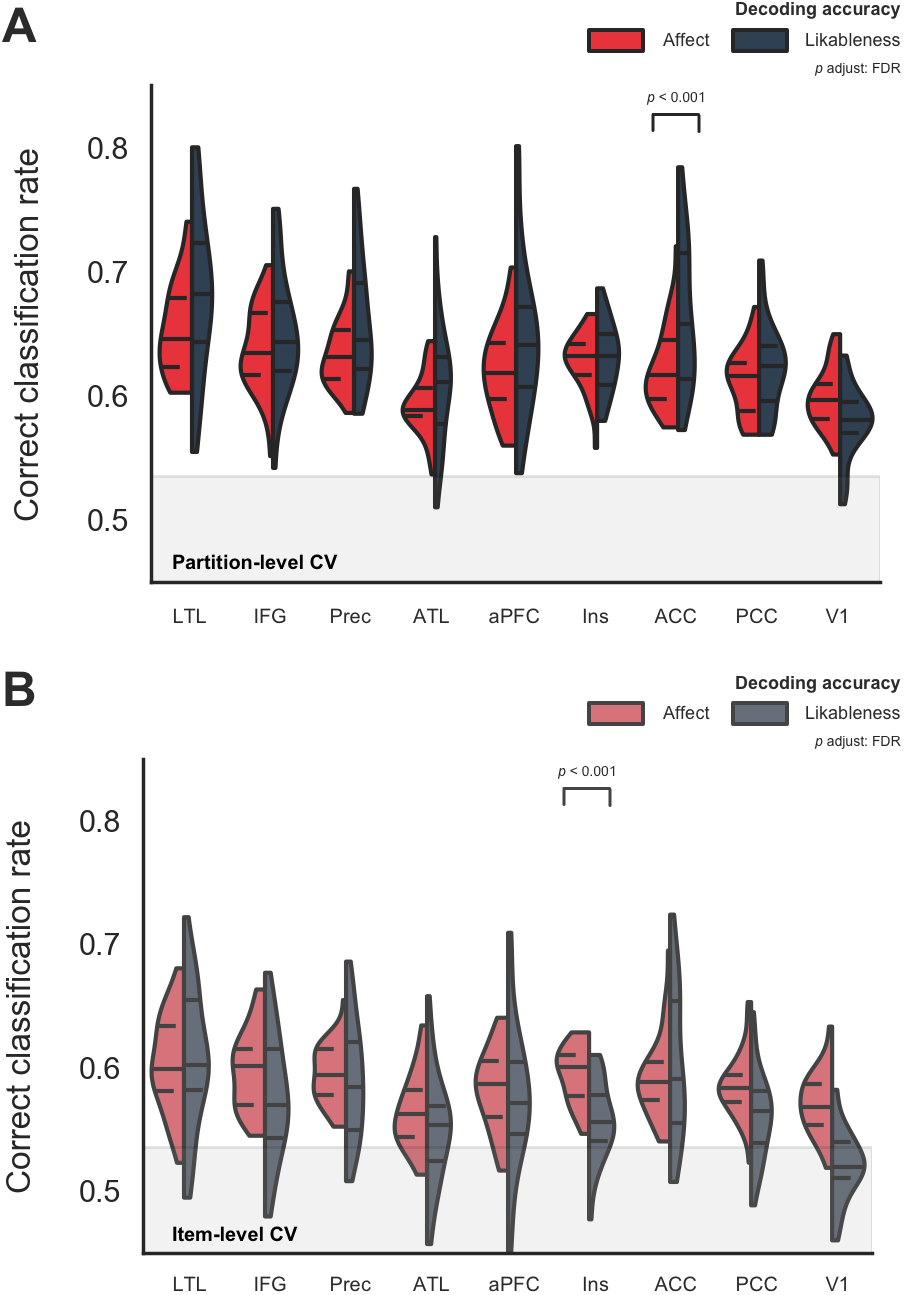
Distributions of decoding accuracies of social concepts. (**A**) Classification accuracy of both the affect and likableness of social knowledge was significantly above chance in all ROIs using a partition-level cross-validation procedure. A repeated-measures ANOVA showed an interaction effect between ROI and concept dimension was driven by the ACC (*p* < .001), which shows a preference for the classification of lik-ableness. The shaded area indicates the average, empirically-estimated chance level (*M* = 0.53). (**B**) Same information but using an item-level cross-validation procedure. Note that the interaction effect between ROI and dimension was driven by the Ins (*p* < .001) instead of the ACC when using the latter cross-validation procedure, showing a preference for the classification of affect.

**Table 5.**
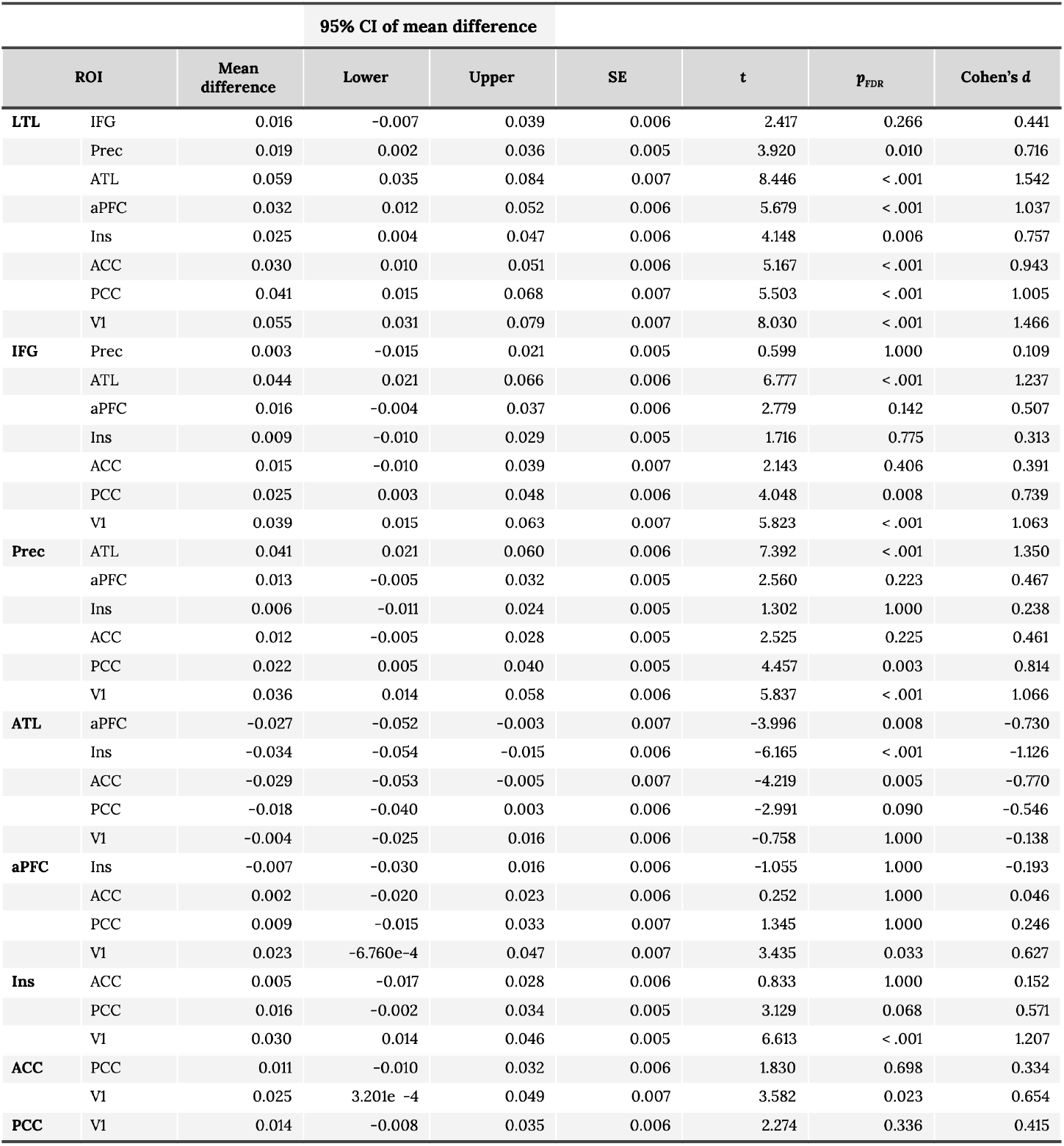
*Post hoc* comparisons of classification accuracies in each pair of ROIs for the main effect of ROI in the binomial classification of affect (high vs. low) using a partition-level cross-validation.

**Table 6.**
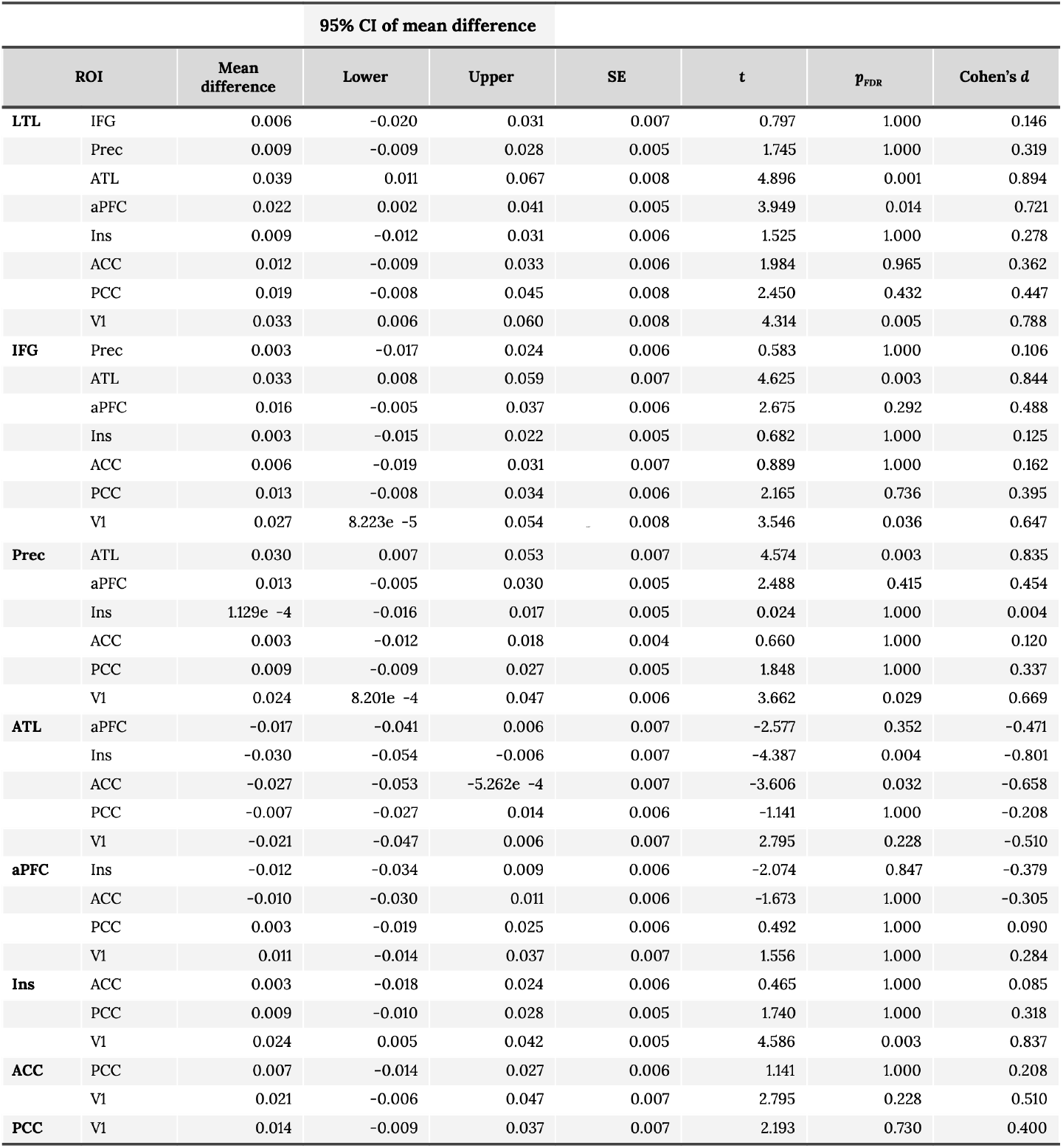
*Post hoc* comparisons of classification accuracies in each pair of ROIs for the main effect of ROI in the binomial classification of affect (high vs. low) using item-level cross-validation.

**Table 7.**
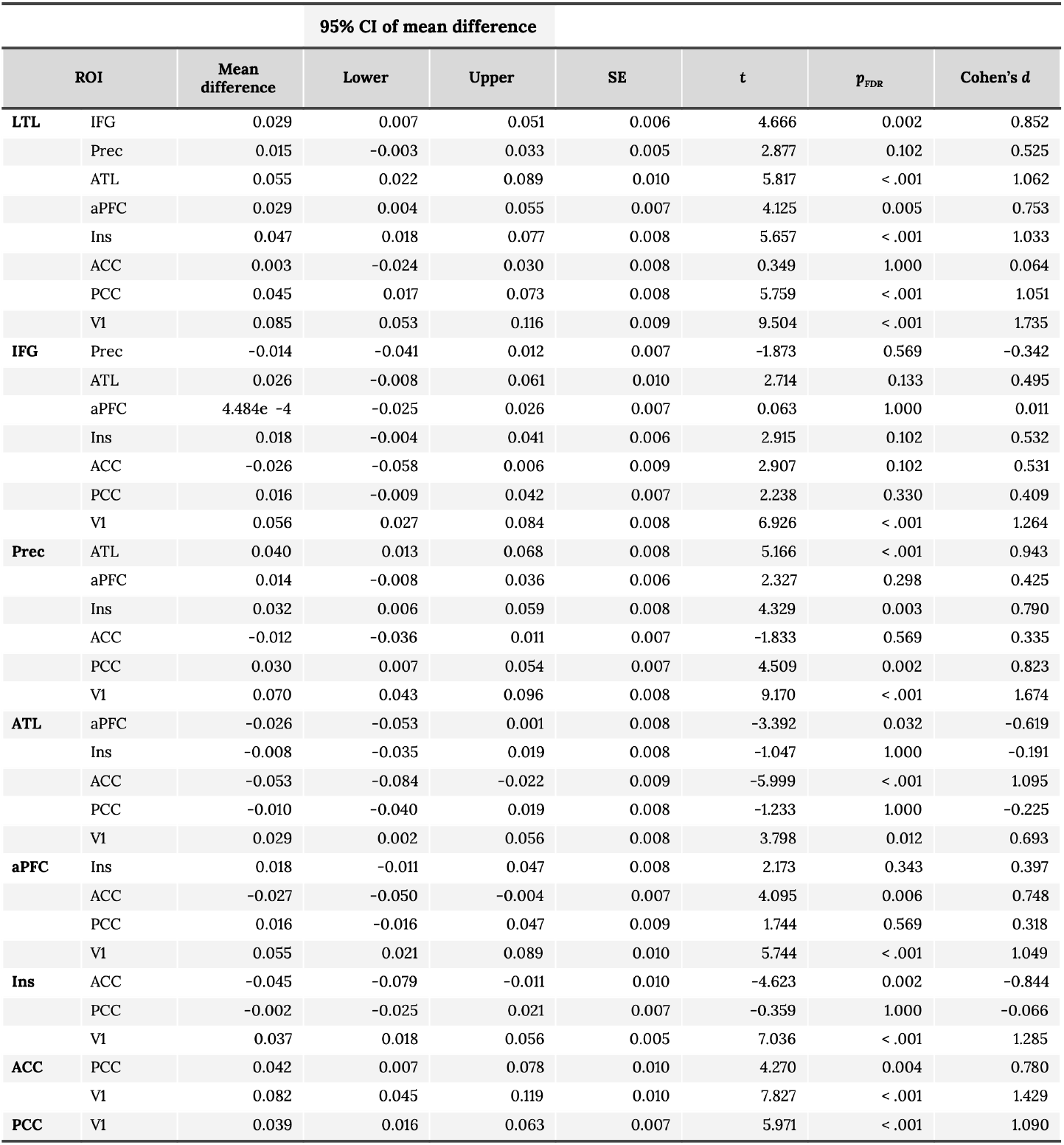
*Post hoc* comparisons of classification accuracies in each pair of ROIs for the main effect of ROI in the binomial classification of likableness (high vs. low) using item-level cross-validation.

**Table 8.** Comparison between the classification accuracies of the affect and likableness of social knowledge. Correct classification rates and summary statistics for the contrast between the classification of likableness (high vs. low) vs. affect (high vs. likableness) using an item-level cross-validation.

| ROI | Likableness | Affect | Mean difference | 95% CI of mean difference |  | SE | t | p<sub>FDR</sub> | Cohen's d |
| --- | --- | --- | --- | --- | --- | --- | --- | --- | --- |
|  |  |  |  | Lower | Upper |  |  |  |  |
| LTL | 0.606 | 0.604 | 0.002 | -0.027 | 0.031 | 0.014 | 0.148 | 0.883 | 0.027 |
| IFG | 0.577 | 0.598 | -0.021 | -0.046 | 0.004 | 0.012 | -1.744 | 0.207 | -0.318 |
| Prec | 0.591 | 0.595 | -0.004 | -0.025 | 0.018 | 0.011 | -0.354 | 0.817 | -0.065 |
| ATL | 0.551 | 0.565 | -0.014 | -0.036 | 0.008 | 0.011 | -1.332 | 0.347 | -0.243 |
| aPFC | 0.577 | 0.582 | -0.006 | -0.035 | 0.023 | 0.014 | -0.402 | 0.817 | -0.073 |
| Ins | 0.559 | 0.595 | -0.036 | -0.052 | -0.020 | 0.008 | -4.623 | < .001 | -0.844 |
| ACC | 0.603 | 0.592 | 0.011 | -0.017 | 0.040 | 0.014 | 0.831 | 0.619 | 0.152 |
| PCC | 0.561 | 0.585 | -0.024 | -0.042 | -0.007 | 0.008 | -2.915 | 0.021 | -0.532 |
| V1 | 0.521 | 0.571 | -0.050 | -0.066 | -0.034 | 0.008 | -6.310 | < .001 | -1.152 |

**Table 9.** Decoding accuracies across concept dimensions. An SVC trained to discriminate the affect dimension (high vs. low) did not generalized to classify examples in the likableness condition, and vice versa, using the item-level CV.

| ROI | Partition-level CV |  | Item-level CV |  |
| --- | --- | --- | --- | --- |
|  | Train Affect<br>–<br>Test Likableness | Train Likableness<br>–<br>Test Affect | Train Affect<br>–<br>Test Likableness | Train Likableness<br>–<br>Test Affect |
| LTL |  |  | 0.511 | 0.496 |
| IFG |  |  | 0.499 | 0.500 |
| Prec |  |  | 0.507 | 0.507 |
| ATL |  |  | 0.490 | 0.499 |
| aPFC |  |  | 0.498 | 0.498 |
| Ins |  |  | 0.496 | 0.492 |
| ACC |  |  | 0.498 | 0.508 |
| PCC |  |  | 0.502 | 0.502 |
| V1 |  |  | 0.508 | 0.508 |

